# Dedicated acetic acid preference coded by broad spectrum ionotropic receptors in a moth species

**DOI:** 10.1101/458646

**Authors:** Rui Tang, Nan-Ji Jiang, Chao Ning, Ling-Qiao Huang, Chen-Zhu Wang

## Abstract

Acetic acid as one of the food related odorant cues attracts many insect species. In the moth *Mythimna separata*, the olfaction of acid was coded via multiple pathways including 3 sensilla types on the antennae and 3 glomeruli in the antennal lobes. Among, suitable dosages of acetic acid exclusively activated DC3 glomerulus that receives integrated projections across sensilla types, which drives attractiveness and feeding attempts of the moth. This circuit encodes broad spectrum ionotropic receptors *8a*, *75q1* and *75q2* which were sufficient to confer acid responsiveness in *Xenopus* oocytes. *Ir75q2* was expressed *in vivo* with *Ir75q1* and it enhanced sensitivity of the receptor functional group toward acids. Furthermore, *Ir75q1* and *Ir75q2* are both necessary for the moth to conduct acetic acid induced reactions of sensilla, DC3 glomerulus as well as attractiveness. Together, it reveals that an indispensable tetramer IR-based unit is employed to fulfill acetic acid specialized preference under suitable dosages through balancing of transcription and peripheral coding. Understanding of the *Ir75q1/2* olfactory pathway provides insights into investigations on acid sensory process in insects.

**Author Summary:** The preference to acids are common in various organisms, and it may involve both olfactory and gustatory reception. In particular, airborne acidity volatiles can be sensed through antennae of insects and later assessed to help locating foraging, mating, and egg laying sites. However, these stimulatory processes can only be delivered by suitable dosages of acids, as we all know that, high acidity could be fatal in most circumstances. To date, avoidance to acids has been well explained in insects, but attractiveness and its basis remain uncharted. In the brain of oriental armyworm *Mythimna separata*, we have located 3 olfactory pathways which may play roles in acetic acid reception. Fortunately, when acetic acid was applied at attractive dosages, it only activated 1 dedicated pathway among the three. Later we found that this attractiveness pathway employed two ionotropic receptor genes namely *Ir75q1* and *Ir75q2*, to successfully deliver this trait. Both genes were necessary for the moth to conduct acetic acid preference, but their roles are different. *Ir75q1* recognized the acetic acid ligand and *Ir75q2* later amplify the sensitivity. By comparing with evidences from electrophysiology and brain imaging tests, we found that the expression bias of either of the two genes has caused the separation of the pathways. It has been revealed in this moth that a smart decision system for olfactory reception exists, and this system may extrapolate to other insect species, as *Ir75q1* and *Ir75q2* are commonly expressed in many insect families.

## Introduction

Insects are precisely developed and possess a complex chemosensory system to communicate with the outside world [1]. Olfaction is critical to this system and conveys crucial information to enable survival in response to multiple chemical cues, including pheromones, host volatiles, or enemy smells [2–4]. Many of these compounds are acids, which exist widely in nature as common products of plants [5,6]. In *Drosophila*, acetic acid is sensed to assess food, oviposition medium, and mating [7–10]. Many other insect species, including armyworms, can be trapped by acetic acid-containing lures [11–15]. However, similar as in humans and nematodes, olfactory delivered acids may also result in opposite behavioral consequences in insects depending on dosage [16,17]. For example, *Drosophila* can avoid high acidity stimuli through specific olfactory receptor neurons (ORNs) which express a family of genes called the ionotropic receptors (IRs) [18,19].

IRs were investigated in variant sensory pathways of *Drosophila* including olfaction [20,21], gustation [22–25], thermosensation [26], and hygrosensation [27,28]. Among, acetic acid is sensed through a spectrum of IRs. The *sensilla coeloconica* on the third chamber of the sacculus express one dedicated *Ir64a*, which mediates repellency toward high acidity stimuli [18]. *Ir64a* is expressed together with a co-receptor *Ir8a* and they together form a dimer-dimer complex to perform as an acid sensing ion channel [20]. Furthermore, *Ir75a* is also used by *D. melanogaster* and *D. sechellia* to smell acetic acid and propionic acid [29,30]. *Ir76b* mediates oviposition preference in female *Drosophila* by sensing acetic acid and citric acid through gustatory reception [10]. Meanwhile, reports on lepidopteran species have also revealed glomeruli based coding for feeding behaviors which were elicited by acidity components [31]. However, the attractiveness olfactory pathways encoded by candidate IR genes to acetic acid itself are still unclear, especially in insects who employ acetic acid as a food related odorant [32].

The Oriental armyworm moth, *Mythimna separata* Walker (Lepidoptera: Noctuidae), also known as *Pseudaletia separata*, is a key pest of crops in Asia and it has been monitored with acetic acid rich lures for decades [33]. In particular, sweet vinegar solution luring is one of the most cost-effective methods of trapping this pest and the recipe for the solution is now a national standard [33]. By changing the proportions of ingredients in such lures, acetic acid-containing lures were found to efficiently trap multiple taxa from different orders, including Lepidoptera [13–15], Diptera [11], Coleoptera [34], and Hymenoptera [12,35]. Discovery of the relevant olfactory perception is essential in both theoretical and application aspects. Due to the high proportion of acetic acid in its luring recipe, we believe that *M. separata* is a suitable target for exploring the olfactory mechanism on acetic acid attractiveness. We hypothesize that acetic acid preference of this moth is mediated by an undiscovered olfactory pathway which contributes to behavioral decision under suitable dosage. As a number of lepidopteran IRs are widely existing across many species [36,37], our results can provide insights to common knowledge on acid driving olfactory and behaviors.

## Results

### *M. separata* antennae respond to acetic acid in the sweet vinegar solution

We first investigated the volatiles from the sweet vinegar solution through the Gas Chromatography coupled with Mass Spectroscopy (GC-MS). Among a total 12 identified chemicals, acetic acid was a major component and covered 7% of the total blend (Table S1). Later in Gas Chromatography coupled with Electroantennographic Detection (GC-EAD), acetic acid evoked dramatic responses in several acetic acid preferred species including *M. separata* (Figure 1A, S1A). However, no response was found in species without an acid preference (Figure S1B). Moreover, reactions of *M. separata* were positively related to acetic acid doses. The sweet vinegar solution which contained equivalent 1000 μg acetic acid has stimulated similar responses as acetic acid solution at the same concentration (Figure 1B). We also found that higher electrophysiological responses were recorded in females other than in males among tested acetic acid dosages (Figure 1B). Therefore, we decided to use only female moths for later experimental process.

**Figure 1.**
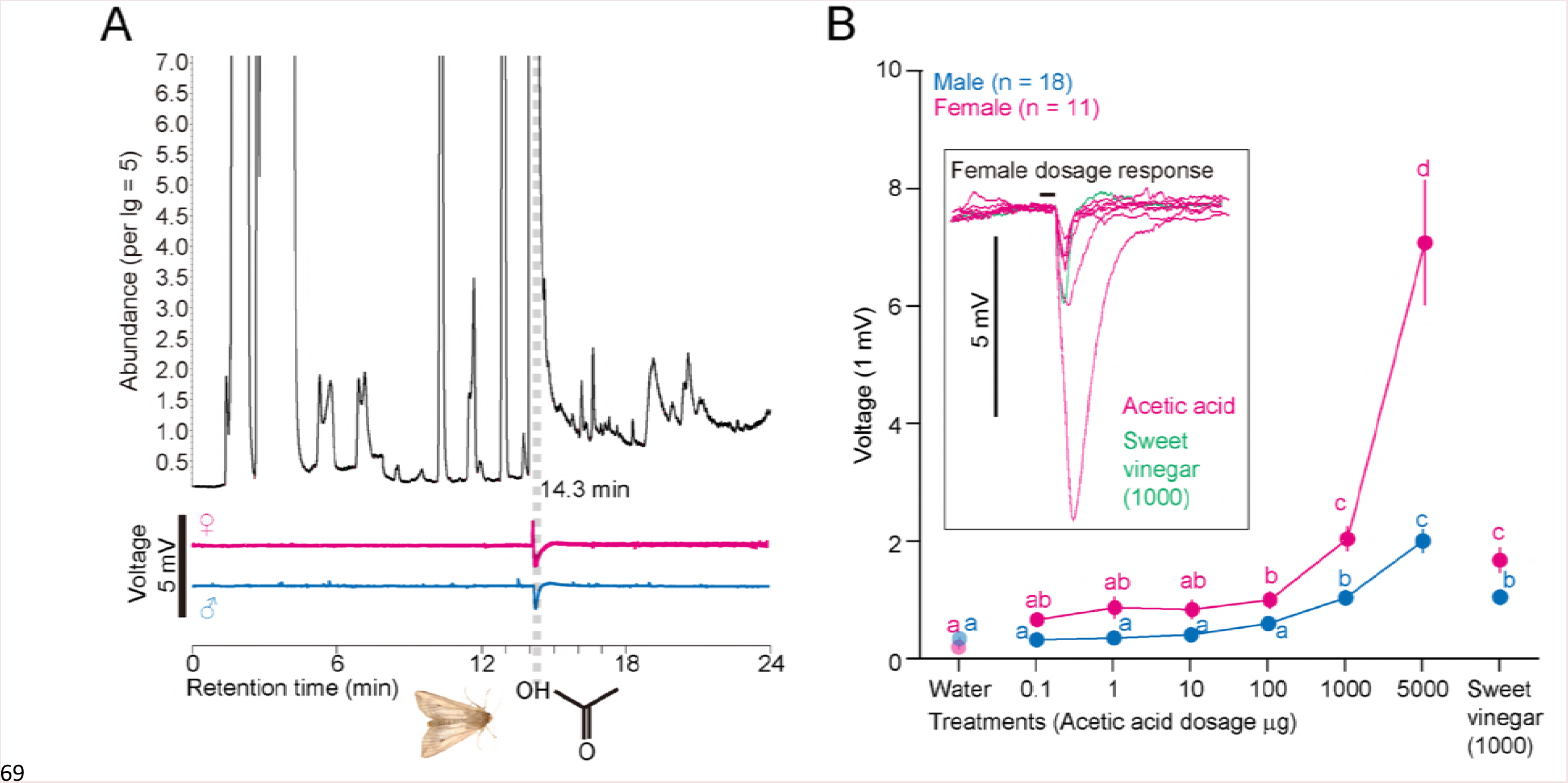
Electrophysiological tests using *M. separata* antennae toward sweet vinegar and laddered acetic acid solutions. **(A)** Example traces showing GC-EAD results with antennae from female (n = 19) and male (n = 4) *M. separata* adults to sweet vinegar solution volatile blends. **(B)** Results from electroantennogram traces and dosage responses with antennae from both genders of *M. separata* toward multiple tested solutions. Water was used as control. The sweet vinegar solution contained equivalent 1000 μg acetic acid as dosage, the formula of the solution was shown in methods section. Amplitudes of female antennae were shown in the square. Different lower case letters indicate significant responses were stimulated among treatments (GLM and Tukey HSD, male: F_7, 138_ = 28.0, *P* < 0.0001, n = 18; female: F_7, 172_ = 63.6, *P* < 0.0001, n = 11). Error bars indicate + s.e.m.

### Three types of sensilla are involved in acetic acid sensing

Next, we conducted single sensillum recordings (SSR) to locate acid sensilla in *M. separata.* Within sensilla from 30 female adults, we identified 38 sensilla with clear responses to acetic acid. Moreover, these sensilla varied in both distribution and firing intensity to certain acids (Figure S2). Spike-count data from all acids were then transferred into z-scores and clustered according to the silhouette method at *k* = 2 to 7 [30,38,39]. The highest silhouette values of 2 to 3 indicated that the sensilla were likely to be classified into 2 or 3 clusters (Figure 2A). Later we found that the sensilla formed three distinguishable areas when plotted with acetic acid, valeric acid and enanthic acid (Figure 2A). A corrgram was used to separate single sensillum into each cluster with manual crosscheck provided (Figure 2B and Data S1). We then named them *as1*, *as2*, and *as3* type sensilla, respectively.

**Figure 2.**
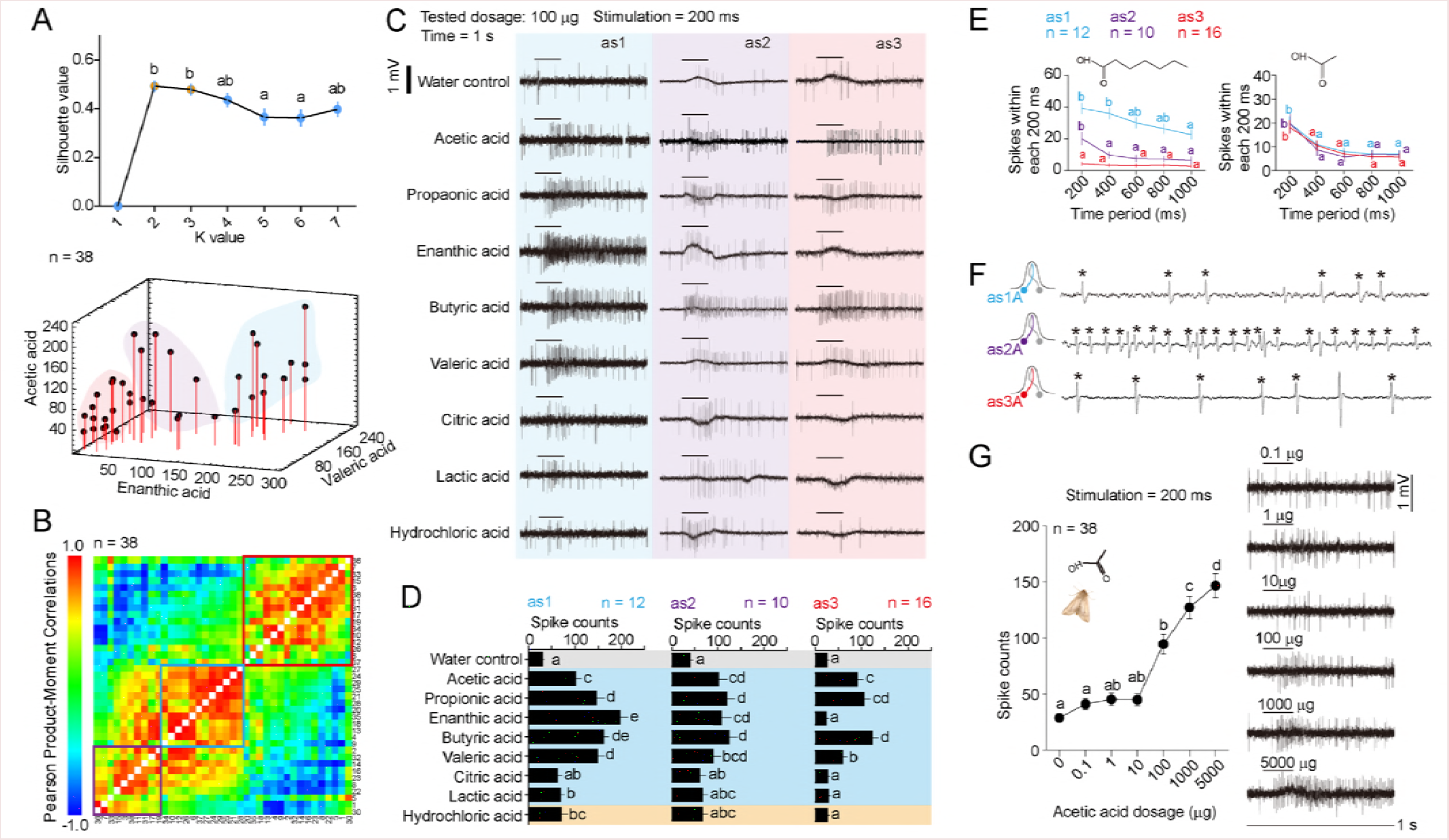
Clustering and characterization of acetic acid active sensilla on female *M. separata* antennae with single sensillum recording (SSR) tests. **(A)** Clustering prediction of sensilla using silhouette method. The peak silhouette value at *k* = 2 and 3 was significantly different from other *k* values (GLM, F_6, 525_ = 42.26, *P* < 0.0001, n = 38). Error bars indicate + s.e.m. Plot of the three acids indicates three areas of clusters. **(B)** Correlation matrix of all sensilla of *M. separata* according to responsiveness patterns of multiple acids. Correlation coefficient was done for each pair of subjects for all tested chemicals including water control, acetic acid, propionic acid, enanthic acid, butyric acid, valeric acid, lactic acid, citric acid, and hydrochloric acid. **(C)** Representative spike patterns of *as1* (n = 12), *as2* (n = 10), and *as3* (n = 16) type sensilla. All tested chemicals were applied at a dosage of 100 μg. Spikes within 1 s were shown. **(D)** Chemical responsiveness patterns of *as1*, *as2*, and *as3* sensilla. Bars with different lower case letters indicate significant differences of spike counts among treatments in each type sensillum (GLM and Tukey HSD, as1: F_8,99_ = 16.98, *P* < 0.0001; as2: F_8,81_ = 3.17, *P* = 0.0036; as3: F_8, 135_ = 16.18, *P* < 0.0001). Error bars indicate + s.e.m. **(E)** Tempo distribution of spikes for enanthic acid and acetic acid in 3 sensilla types. Different lower case letters indicate significant decrease of spikes in each successive 200 ms (GLM and Tukey HSD, enanthic acid, as1: F_4, 55_ = 4.42, *P* = 0.0036, as2: F_4,45_ = 4.84, *P* = 0.0025, as3: F_4, 75_ = 0.28, *P* = 0.89; acetic acid, as1: F_4, 55_ = 7.72, *P* < 0.0001, as2: F_4, 45_ = 5.49, *P* = 0.0011, as3: F_4, 75_ = 7.58, *P* < 0.0001). Enanthic elicited tonic responses in as1 and no response in as3, while other patterns were all phasic. Error bars indicate + s.e.m. **(F)** Spike sorting in *as1, as2*, and *as3* type sensilla. Asterisks indicate acid responding ORNs in each sensilla type, respectively. **(G)** The dose responses of all sensilla pooled to acetic acid. Representative reactions were shown within 1 s. Dots with different lower case letters indicate significant differences of spike counts among dosages (GLM and Tukey HSD, F_6, 259_ = 39.78, *P* < 0.0001). Error bars indicate + s.e.m.

All three types of sensilla responded broadly to several acids with different spike counts (Figure 2C, Data S1). Among, *as1* sensilla significantly responded to 7 of 8 tested acids including acetic acid, propionic acid, enanthic acid, butyric acid, valeric acid, lactic acid, and hydrochloric acid. While *as2* responded to 5 of them and *as3* responded to only 4 (Figure 2C, 2D). Furthermore, the tempo distribution of spikes varied among sensilla types for enanthic acid. A moderate tonic reaction pattern was found in *as1*, yet phasic patterns were observed in other types. While acetic acid stimulated only phasic pattern across sensilla (Figure 2E). Multiple amplitudes were found in all sensilla types, indicating that numbers of ORNs were housed (Figure 2C, 2F). By manually sorting and comparing spikes to all active acids, we suggested that only one ORN was activated in each sensilla type (Figure 2F, Data S1). Since acetic acid elicited similar spike counts with undistinguished tempo-spacial distributions, we then pooled the dosage responses for all ORNs. Significant reactions could be observed when applied with 100 μg acetic acid, and an increase was found along with dosages (Figure 2G).

### Acetic acid stimulates attractiveness and feeding attempt in *M. separata* in a dosage manner

In wind tunnel tests, both the sweet vinegar solution and 10 - 1000 μg acetic acids elicited comparable attractiveness to *M. separata*, excepted for landing behavior (Figure 3C). Dosages of 10 μg to 1000 μg acetic acids all caused significant upwind flights comparing to the water control. Moreover, we found that 12 h fasted moths adults conducted significant attractiveness behaviors to acetic acid than fed moths (Figure 3B, 3C). We asked whether this feature is specific for acetic acid by comparing it to two non-food related odorant acid chemicals propionic acid (related to acetic acid in *Drosophila* IR sensory) and enanthic acid (performed very well in SSR tests). Interestingly, the result revealed that acetic acid was the only one which elicited significant upwind flights to the moth among acids (Figure 3C and Figure S3). Furthermore, an overwhelming of 5000 μg acetic acid terminated attractiveness in the moths.

**Figure 3.**
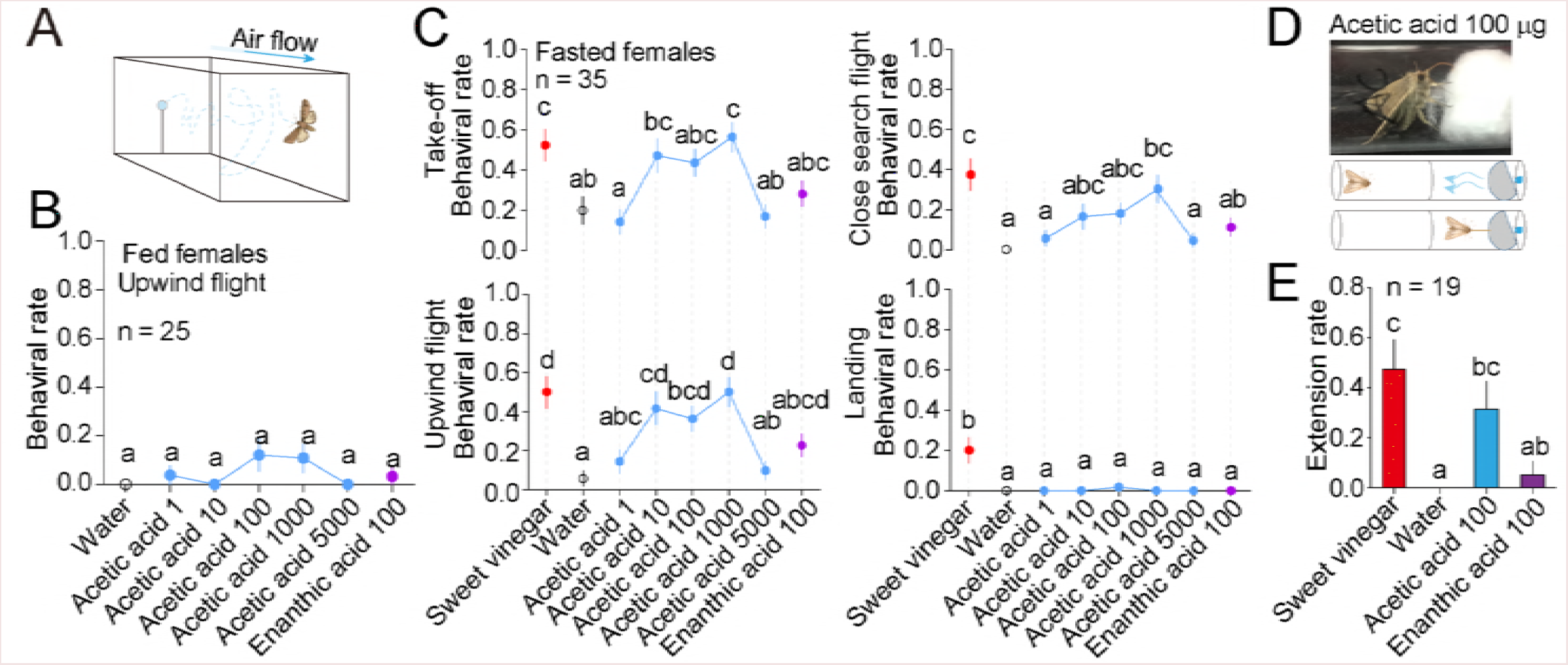
Behavioral responses of *M. separata* in indoor assays. The dosage unit in all tests was μg. The sweet vinegar solution was used at equivalent dosages as 1000 μg acetic acid. **(A)** Schematic diagram shows wind tunnel layouts. **(B)** Comparison of upwind flight behaviors elicited by multiple acidity odorant sources in fed female adults. No difference was observed among treatments (GLM and Tukey HSD, F_6, 187_ = 1.8, *P* = 0.97). Error bars indicate + s.e.m. **(C)** Comparison of four criterial behaviors elicited by multiple acidity odorant sources in 12 h fasted female moths. Different lower case letters indicate significant differences among treatments (GLM and Tukey HSD, take-off: F_7, 333_ = 5.5, *P* < 0.0001; upwind flight: F_7, 333_ = 6.9, *P* < 0.0001; close search: F_7, 333_ = 5.4, *P* < 0.0001; landing: F_7_, _333_ = 8.9, *P* < 0.0001). Error bars indicate + s.e.m. **(D)** Schematic diagram shows PER assay layouts, blue drops indicate tested odorants which were separated from insects to avoid gustatory reception. **(E)** Comparison of proboscis extension rates elicited by multiple acidity odorant sources. Different lower case letters indicate significant differences among treatments (GLM and Tukey HSD, F_3, 72_ = 7.0, *P* < 0.0001). Error bars indicate + s.e.m.

As feeding status significantly influenced behavioral outputs, we then tested the fasted moths in an adopted proboscis extension response (PER) assay with olfactory stimuli [40] (Figure 3D). Results showed that sweet vinegar solution and acetic acid both elicited significant PER behaviors comparing to water control (Figure 3E). This clearly revealed that acetic acid can serve both at range and closely as a food related odorant source (Movie S1). On the other, enanthic acid again has hardly initiated PERs, indicating different olfactory signaling exists according to acid types.

### DC3 glomerulus is exclusively evoked by acetic acid under attractive dosages

We then traced the acetic acid signaling in the antennal lobe, the first olfactory neuropil of the insect’s brain. In *in vivo* optical imaging tests, a total three different regions of interest (ROIs) were evoked by multiple acidity components (Figure 4A). A pattern change was observed in acetic acid stimuli: only 1 area was constantly evoked during the dosages of 10 to 1000 μg, while 3 areas were evoked when applied at 5000 μg (Movie S2). Enanthic acid evoked only 1 area with weak intensity regardless of dosages (Movie S3). Propionic acid, butyric acid, and valeric acid all evoked 3 areas of interests (Figure 4A). Other tested acids did not show a significant evoking patterns under the scope of this study (Figure 4C and Figure S4).

**Figure 4.**
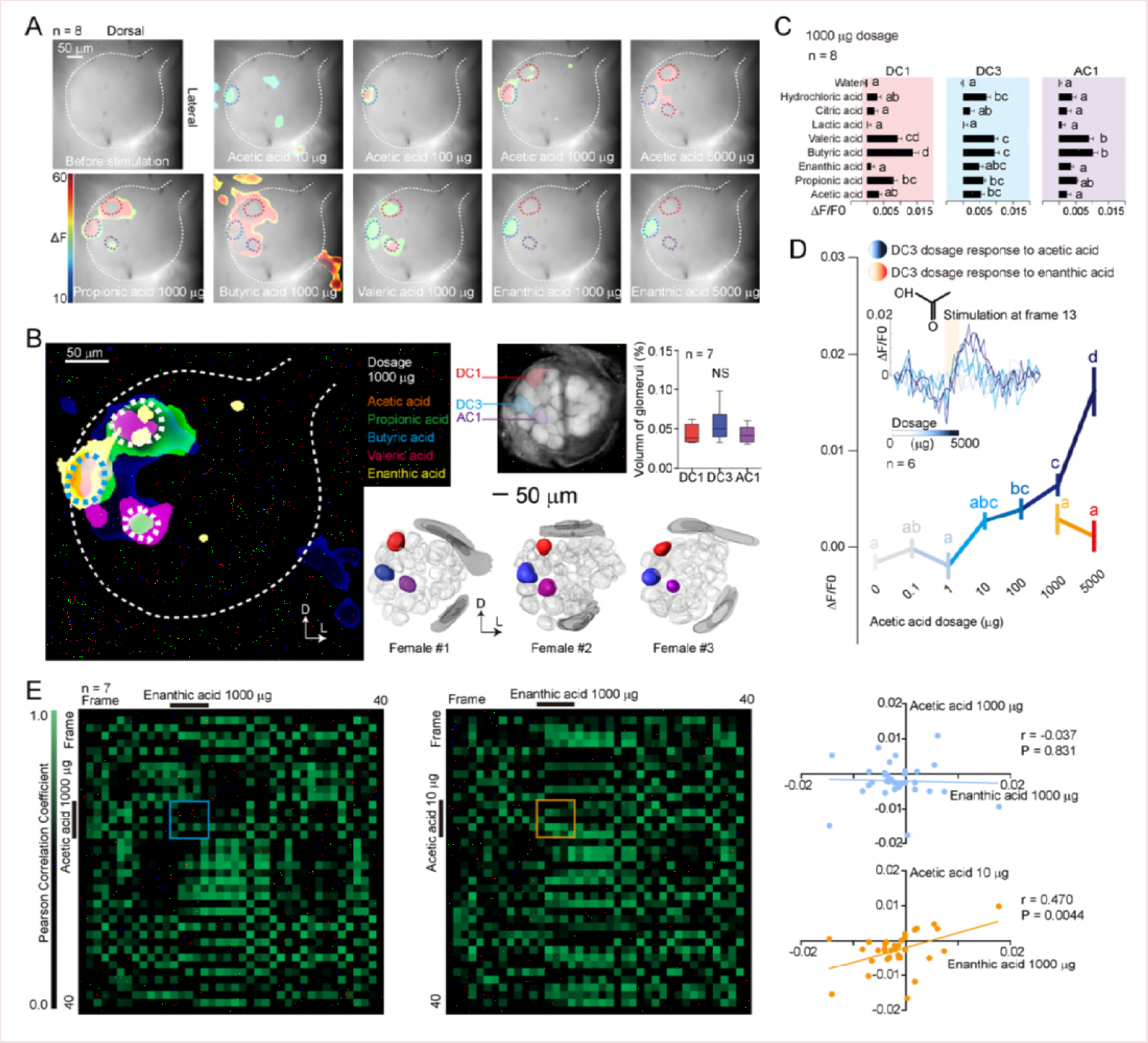
Representative antennal lobe optical imaging patterns and glomeruli identification of female *M. separata* adults to acids. **(A)** Representative evoking patterns in *M. separata* female antennal lobes across chemicals & dosages. Color circles indicate areas of interests. **(B)** Localization and naming of glomeruli which can be evoked by acetic acid. Left: merged evoking areas from top 5 active acids. Right: brain atlas of female adults. The three glomeruli were named according to locations as DC1 (dorsal central 1), DC3, and AC1 (anterior central 1), respectively. Bar chart shows volume statistics in three glomeruli (GLM and Tukey HSD, F_2, 18_ = 1.26, *P* = 0.308). Error bars indicate + s.e.m. **(C)** Comparison of evoking patterns among acids in three glomeruli. Bars with different lower case letters indicate significant differences of *ΔF/F0* values among treatments (GLM and Tukey HSD, DC1: F_8,198_ = 19.7, *P* < 0.0001; DC3: F_8, 198_ = 10.4, *P* < 0.0001; AC1: F_8, 198_ = 10.7, *P* < 0.0001). Error bars indicate + s.e.m. **(D)** Dosage response tests of DC3 glomerulus to acetic acid as it was the only active glomerulus under attractive dosages. Dots with different lower case letters indicate significant differences of *ΔF/F0* values among dosages (GLM and Tukey HSD, acetic acid: F_5, 102_ = 24.7, *P* < 0.0001; enanthic acid: *P* = 0.377). Error bars indicate + s.e.m. **(E)** Similarity analysis in antennal lobe reaction patterns among acids by cross correlations. Reactions were taken from DC3 area for each acid and reactions were compared in a pair wise manner along 40 frames. Green areas indicate highly correlated evoking patterns between the 2 tested acids. Squares indicate glomeruli reactions during stimulation time. No significant correlation was observed between reactions of 1000 μg enanthic acid and 1000 μg acetic acid (Pearson’s correlation, *r* = −0.037, *P* = 0.831), a shallow but significant correlation was observed between reactions of 1000 μg enanthic acid and 10 μg acetic acid (Pearson’s correlation, *r* = 0.470, *P* = 0.0044).

To identify correspondence glomeruli, we merged ROIs from the above 5 active acids. Result showed that the ROIs from all acids were highly overlapped, indicating 1 glomerulus was involved in each area (Figure 4B). We later established 3-D structures of *M. separata* female antennal lobes with a standard atlas protocol [41–43], and compared the morphology of glomeruli with imaging results. The glomeruli reflecting ROIs were identical in terms of positions among individuals and no significant difference was observed in volumes. We then named them according to locations as DC1, DC3, and AC1 (Figure 4B). When all acids were tested at 1000 μg, DC1 responded to only propionic acid, butyric acid, and valeric acid. AC1 responded only to butyric acid and valeric acid. Most importantly, DC3 was the only glomerulus among the three with acetic acid activity under this dosage, and this glomerulus had the broadest recognition spectrum towards acids (Figure 4C). Reactions of DC3 also started at a dosage of 100 μg and it increased with dosages of acetic acid, but this glomerulus did not respond to increasing of enanthic acid dosage (Figure 4D). It was obvious that DC3 received integrated axon projections from multiple sensilla types as per the SSR test results.

To better address the relationships of antennal lobe evoking patterns with behavioral consequences, we cross-correlated these acids in a pair wise comparison along 40 frames in DC3 areas (Figure 4E). During stimuli, results showed that 1000 μg acetic acid did not correlate well with 1000 μg enanthic acid (blue square, Figure 4E), which is consistent with different behavioral outputs in wind tunnel tests for these acids. On the other, a shallow yet significant correlation was observed between 10 μg acetic acid and 1000 μg enanthic acid, indicating that evoking intensity of DC3 has contributed in close search behaviors (orange square, Figure 4E). To sum up, so far evidences showed that antennal lobe reaction patterns could be the main driving force for determination of behavioral outputs to acids, and DC3 is highly likely to be the main glomerulus involved in acetic acid stimulated attractiveness during 100 to 1000 μg dosages. With results have been obtained on behavioral and cellular basis of acetic acid response, we started to examine its molecular basis.

### Putative acetic acid sensing receptors *Ir75q1*, *Ir75q2* and *Ir8* are co-expressed *in vivo*

We then obtained the antennal transcriptome of *M. separata* adults and screened for putative receptors with BLAST against acid sensing IRs of *Drosophila melanogaster*. A total 6 IRs were found and named as *Ir8a*, *Ir75q1*, *Ir75q2*, *Ir64a*, *Ir75d*, and *Ir75p*, respectively (Figure 5A). However, after rounds of PCR tests (both ORF cloning and reverse transcription PCR), we only managed to obtain consistent CDSs of 4 genes - *Ir8a*, *Ir75q1*, *Ir75q2*, and *Ir64a*. The tissue-gender expression levels of the 4 genes were similar with results from transcriptome analysis (Figure 5A and Figure S5). Among, *Ir8a* was abundantly expressed in the antennae, indicating its co-receptor role in IR-based olfaction [20]. While *Ir64a* was the homologous with *Drosophila Ir64a* with the highest similarities revealed in both amino acid alignments and initial structural prediction, indicating its potential hydrogen ion sensory activity (Figure S7). As our purpose was to identify the acetic acid attractive pathway, we then emphasized on the left *Ir75q1* and *Ir75q2*.

**Figure 5.**
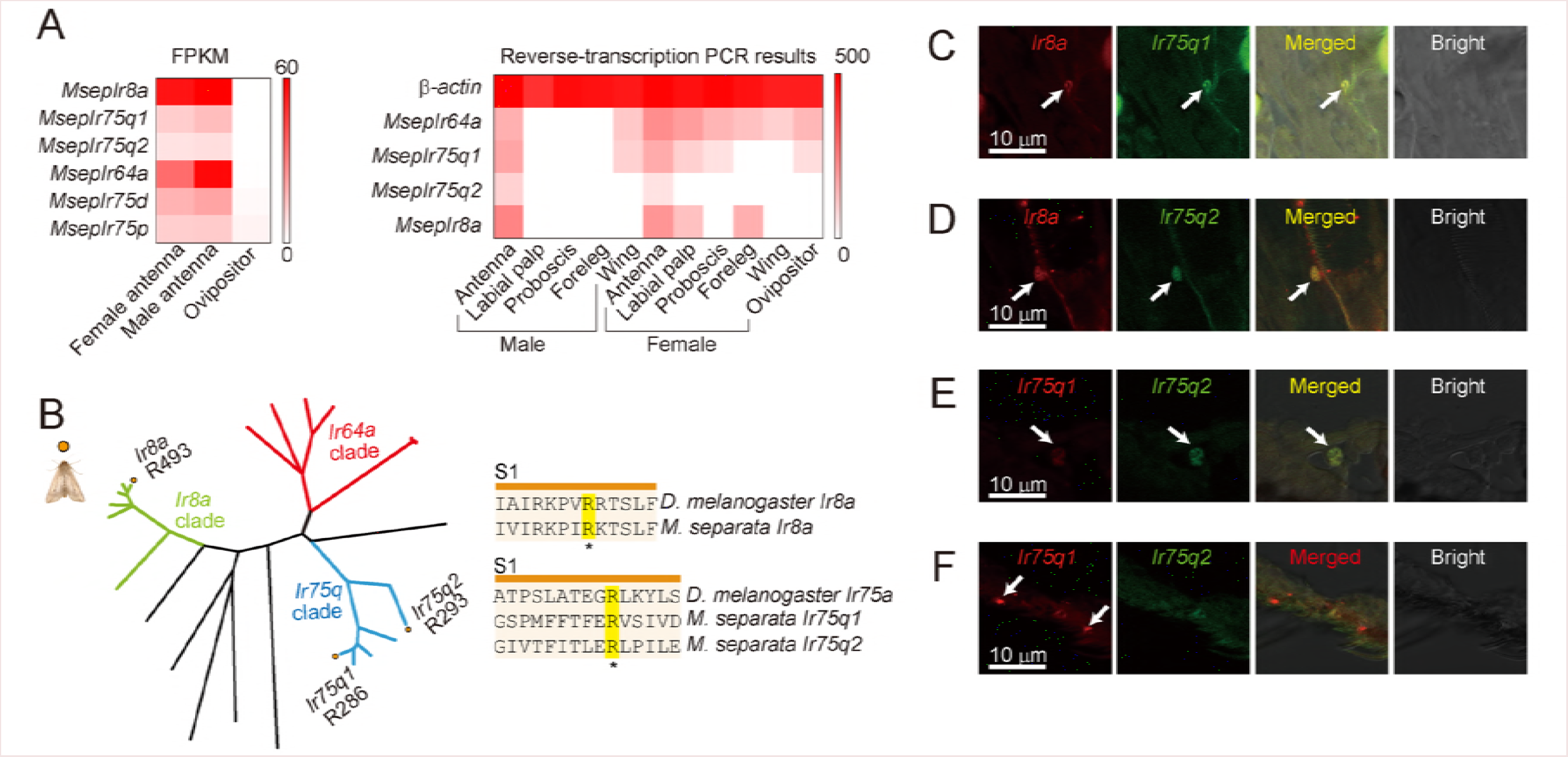
Screening of putative acetic acid sensing IRs in *M. separata*. **(A)** Tissue expression of IRs by RNA-seq and reverse transcription PCR (RT-PCR) using cDNAs from different body parts of *M. separata*. *β*-actin was used as reference. **(B)** Phylogenetic analysis of related IRs from multiple species. Deposited IR gene data and protocol were shown in Table S3. *: predicted carboxyl recognition arginine residues. **(C-F)** Co-expression patterns of *Ir8a*, *Ir75q1*, and *Ir75q2* in *M. separata* female antennae. Arrows show labelled somata with probes synthesized from targeted genes. Digoxin (red) and biotin (green) were used to label the two genes respectively in each pair.

*Ir8a* had a dimer structure, while both *Ir75q1* and *Ir75q2* had a monomeric structure, which was similar to *D. melanogaster* acid olfactory couterparts (Figure S6 and Figure S7). By comparing with structural predictions of vinegar fly IRs [19], a conserved arginine (R) residue was observed in S1 region of ligand binding domain for all three receptors, indicating potential carboxyl binding activity of all IRs (Figure 5B and Figure S6). *Ir8a* was structurally more conserved with ionotropic glutamate receptors (iGluRs) as it maintains the conserved aspartate (D) in S2 region. However, the corresponding residues of *Ir75q1* and *Ir75q2* were shifted, which means a functional change may occur for these IRs from glutamate binding activity to acid (Figure 5B and Figure S6).

One thing special was that, *Ir75q2* (FPKM = 7.638, females) was always found together with *Ir75q1* (FPKM = 15.045, females) in the same assembled contig. We, therefore, speculated that the functions of *Ir75q1* and *Ir75q2* were closely related and they were likely to be expressed in the same ORNs for *M. separata* to accelerate acid detection. To investigate this, we conducted two-color *in situ* hybridization with probes for these four genes. The results showed that both *Ir75q1* and *Ir75q2* were consistently co-expressed with *Ir8a* in the antennae of female *M. separata* adults (Figure 5C, 5D), which reveals that *Ir8a* serves as a co-receptor in acid reception. As predicted, *Ir75q2* was always co-expressed with *Ir75q1* (Figure 5E), yet single expression of *Ir75q1* could also be observed (Figure 5F).

### Functional architectures of *Ir75q1* and *Ir75q2*

We investigated *Ir8a*, *Ir75q1*, and *Ir75q2* with *Xenopus* oocyte expression system which allows us to assemble every possible combination of interested IRs. The cells were challenged with multiple acid solutions at the concentration of 10^−6^ mol/L to avoid significantly changing to the pH (Figure S9A).

Among all treatments, *Ir8a* and *Ir75q1* injected oocytes responded to all acids at low rates and an acetic acid preference was observed. While *Ir8a* and *Ir75q2* injected cells had a diagnostic response to enanthic acid. When *Ir75q1*, *Ir75q2* and *Ir8a* were injected together into one cell, still there was a high response to enanthic acid, but the responses to all acids including acetic acid were significantly amplified (Figure 6A). Meanwhile, a high sensitivity to acetic acid was observed for *Ir8a/75q1/75q2* cells, with plateau at concentrations ≤ 10^−6^ mol/L (Figure 6B). These results indicate that the *Ir8a/75q1/75q2* group may serve in low acetic acid sensing pathways. Moreover, for *Ir8a/75q1/75q2* expressing cells, all acids stimulated responsiveness regardless of pH levels of tested solutions (Figure 6C and Figure S9). This suggests that *Ir75q1/q2/8a* serves as a ligand-based channel other than pH-based.

**Figure 6.**
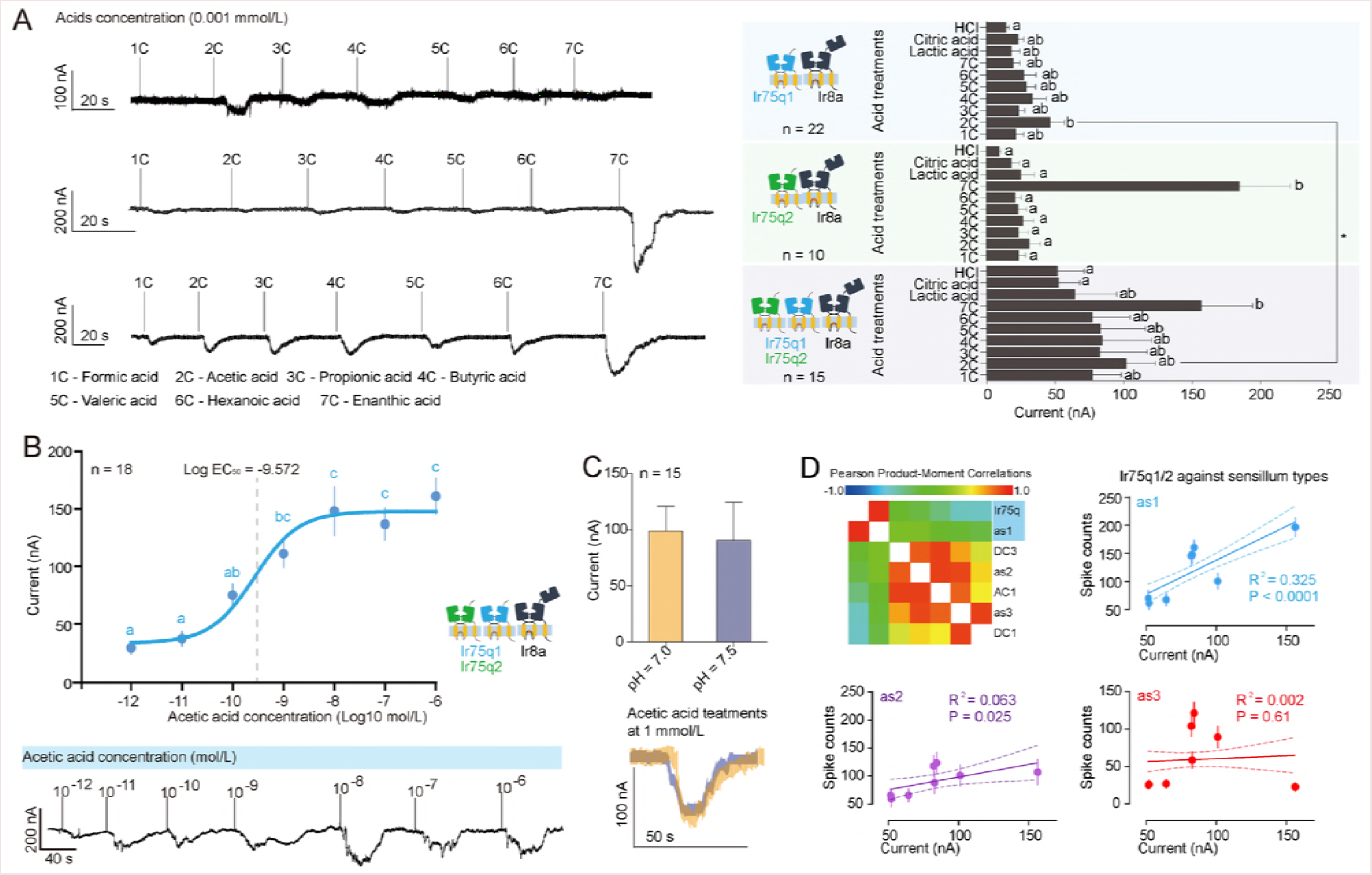
Functional characterizations of *Ir8a*, *Ir75q1*, and *Ir75q2* using *Xenopus* oocyte expression system. **(A)** Representative responses and statistics of cells tested with multiple acid solutions. 1C: formic acid, 2C: acetic acid, 3C: propionic acid, 4C: butyric acid, 5C: valeric acid, 6C: hexanoic acid, 7C: enanthic acid. Bars with different lower case letters indicate significant differences of responses caused among tested acids (GLM and Tukey HSD, *Ir8a/Ir75q1*: F_9, 278_ = 1.9, *P* = 0.052; *Ir8a/Ir75q2*: F_9, 100_ = 5.97, *P* < 0.0001; *Ir8a/Ir75q1/Ir75q2*: F_9, 219_ = 2.36, *P* = 0.014). *: significant higher responses were caused by *Ir8a/Ir75q1/Ir75q2* compared with *Ir8a/Ir75q1* (GLM, F_90_ = 14.94, *P* < 0.0001). Error bars indicate + s.e.m. **(B)** Dosage responses of *Ir8a/Ir75q1/Ir75q2* to laddered acetic acid solutions. Dots with different lower case letters indicate significant differences of responses caused among concentrations (GLM and Tukey HSD, F_6, 135_ = 14.9, *P* < 0.0001). Mean Log EC_50_ = −9.572 was shown. Error bars indicate + s.e.m. **(C)** Acidity responses of *Ir8a/Ir75q1/Ir75q2* to normal (pH 7.0) and neutralized (pH 7.5) acetic acid solutions with the same concentration of 10^−3^ mol/L. Error bars indicate + s.e.m. **(D)** Correlation matrix for IRs, sensilla and glomeruli according to multiple responsiveness patterns to acids. Dot charts show correlations of acids stimulated reaction patterns between *Ir8a/Ir75q1/Ir75q2* and each type sensilla, respectively. 95% confidence interval was shown between dotted lines in each chart.

For all tested acid chemicals, no observable response was recorded with the intact oocyte control, as well as treatments of *Ir8a*, *Ir75q1*, or *Ir75q2* alone injection (Figure S8A-D, S8G). Meanwhile, reaction patterns to multiple acids of *Ir75q1/q2/8a* perfectly coincided with as1A, and a shallow yet significant correlation with as2A (Figure 6D).

### *Ir75q1* and *Ir75q2* are necessary for *M. separata* to conduct acetic acid attractiveness behavior

An RNA interference (RNAi) test was carried out to further investigate the necessity of *Ir75q1* and *Ir75q2* in acetic acid attractiveness of *M. separata*. By injecting small interfering RNAs (siRNAs) of both *Ir75q1* and *Ir75q2*, we observed that expression levels of both genes were significantly down-regulated (Figure 7A). Knock-down of *Ir75q1* and *Ir75q2* resulted in several consequences. Firstly, spike counts of acid sensory neurons were significantly decreased for acetic acid and enanthic acid in siIr75q1/2 moths (Figure 7C and Data S2). Secondly, siIr75q1/2 moths showed a significantly decreased DC3 overlapped area activity to both acetic acid and enanthic acid (Movie S4). On the other hand, DC1 area activity was hardly influenced. There was a trend of decrease in AC1 area activity yet the effect was not significant (GLM, *P* = 0. 224), indicating possible involvement of *Ir75q1* within this circuit (Figure 7D). Last, siIr75q1/2 moths showed significantly decreased attractiveness behaviors in wind tunnel tests to both acetic acid and sweet vinegar solution (Figure 7B). siRNAs from the green fluorescent (GFP) protein were used to assess off-target effect. Injection of siGFPs did not cause down-regulation of either *Ir75q1* or *Ir75q2* (Figure 7A). Attractiveness of siGFP moths to acetic acid and sweet vinegar solution was not significantly influenced in wind tunnel (Figure 7B). The results from RNAi tests not only confirmed the existence of *Ir75q1/2-as-DC3* pathway but also clearly indicated that this pathway acts as an indispensable stimulatory olfactory pathway which drives acetic acid attractiveness behaviors in *M. separata* adults.

**Figure 7.**
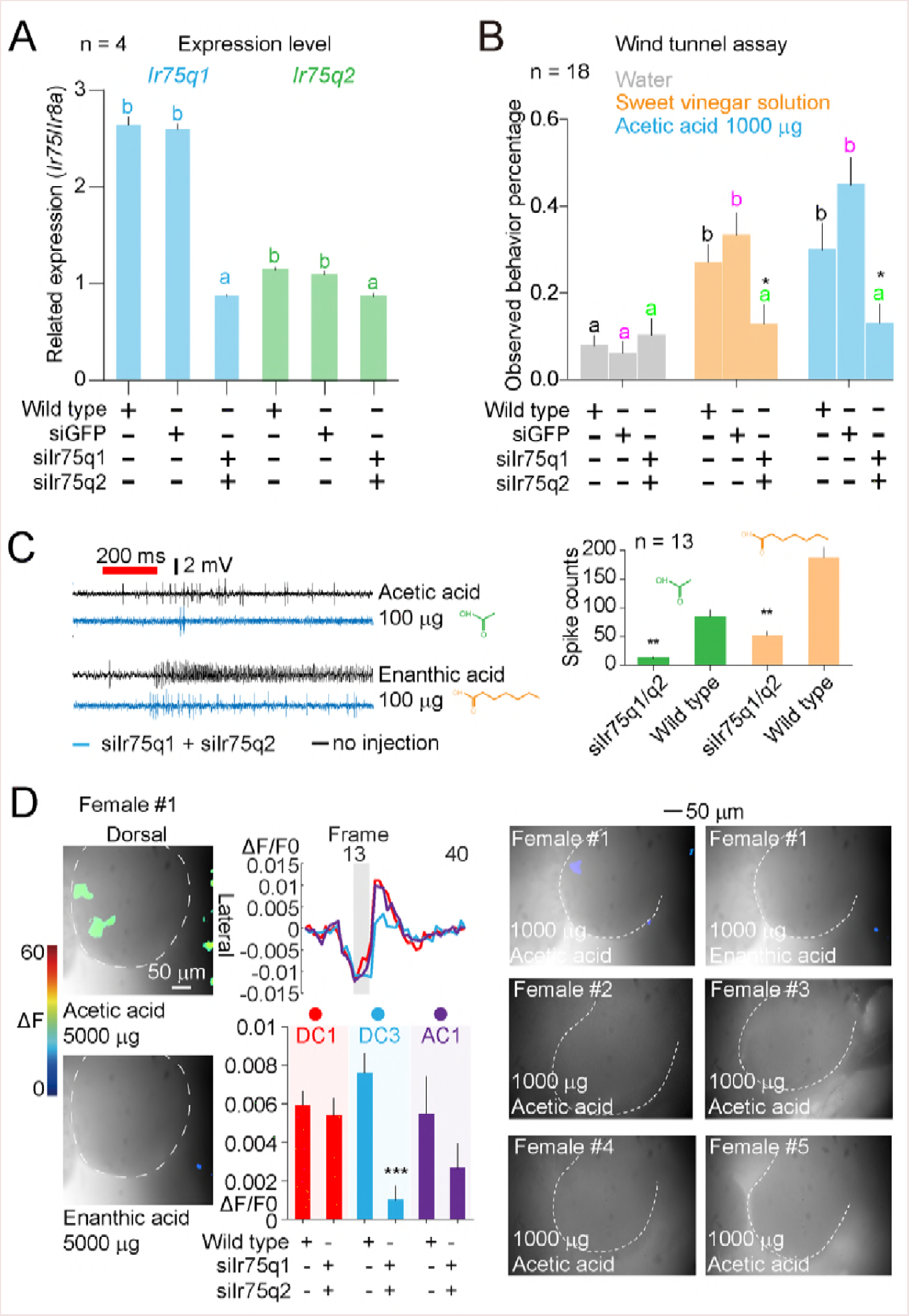
Results of RNAi tests using siRNAs from *Ir75q1* and *Ir75q2*. **(A)** Comparison of *Ir75q* gene expression levels between siIr75q strain and control strain. Expression level was calculated according to 1/power (2, *Ir75q* - *Ir8a)*. Off-target effect was assessed by injection of siGFP RNAs. n = 4 with 3 technical replicates in each data unit. Different lower case letters indicate significant lower expression levels of *Ir75q1* and *Ir75q2* than wild type strain (GLM and Tukey HSD, *Ir75q1*: F_2, 9_ = 295, *P* < 0.0001; *Ir75q2*: F_2, 9_ = 19.1, *P* < 0.0001). Error bars indicate + s.e.m. **(B)** Wind tunnel assay among siIr75q strain, siGFP strain, and wild type strain. Bars with different lower case letters with the same color indicate significant differences of observed positive behavior percentages among treatments in each strain (GLM and Tukey HSD, siGFP strain: *F_2,54_* = 16.4, *P* < 0.0001; wild type: F_2, 108_ = 8.5, *P* = 0.0004.) Error bars indicate + s.e.m. Bars with the same lower case letters in green indicate no significant difference was observed in positive behavior percentages among treatments in siIr75q injected strain (GLM and Tukey HSD, F_2,61_ = 0.14, *P* = 0.867). * significant differences were observed between siIr75q strain and wild type strain to the same odorant treatment (Student’s *t* test, water: *t* = 0.892, *P* = 0.378; acetic acid 1000 μg: *t* = 2.16, *P* = 0.036; sweet vinegar solution: *t* = 2.04, *P* = 0.046). No difference was observed between siGFP injected strain and control strain to the same odorant treatment (Student’s *t* test, water: *t* = 0.466, *P* = 0.64; acetic acid 1000 μg: *t* = 0.891, *P* = 0.38; sweet vinegar solution: *t* = 1.63, *P* = 0.11). **(C)** Comparison of spike counts from acid sensilla to acetic acid and enanthic acid between siIr75q strain and wild type strain. ** significant decrease of spike counts of siIr75q strain than wild type strain (Student’s *t* test, acetic acid: *t* = 4.8, *P* < 0.0001; enanthic acid: *t* = 6.6, *P* < 0.0001). Error bars indicate + s.e.m. **(D)** Comparison of glomeruli activities elicited by acetic acid and enanthic acid between siIr75q strain and wild type strain. Bar chart indicate evoking intensities sampled from anticipated glomeruli areas unless observable reactions were seen. ** significant decrease of activity of siIr75q DC3 glomerulus than wild type strain (Student’s *t* test, DC1: *t* = 0.65, *P* = 0.519; DC3: *t* = 3.7, *P* = 0.0008; AC1: *t* = 0.13, *P* = 0.900). Error bars indicate + s.e.m. Right: replicates showing termination of glomeruli reactions to acetic acid and enanthic acid at a dosage of 1000 μg.

We then established a schematic to show sensing mechanism of acids in *M. separata* (Figure 8A, 8B). To assess this model, we knocked-down each gene one by one and tested moth adults with an EAG approach. Results showed that, single injection of siIr8a, siIr75q1, or siIr75q2 all significantly decreased reactions to acetic acid (Figure 8C). While for enanthic acid, only siIr75q2 strain showed a significant decrease compared to wild type (Figure 8C).

**Figure 8.**
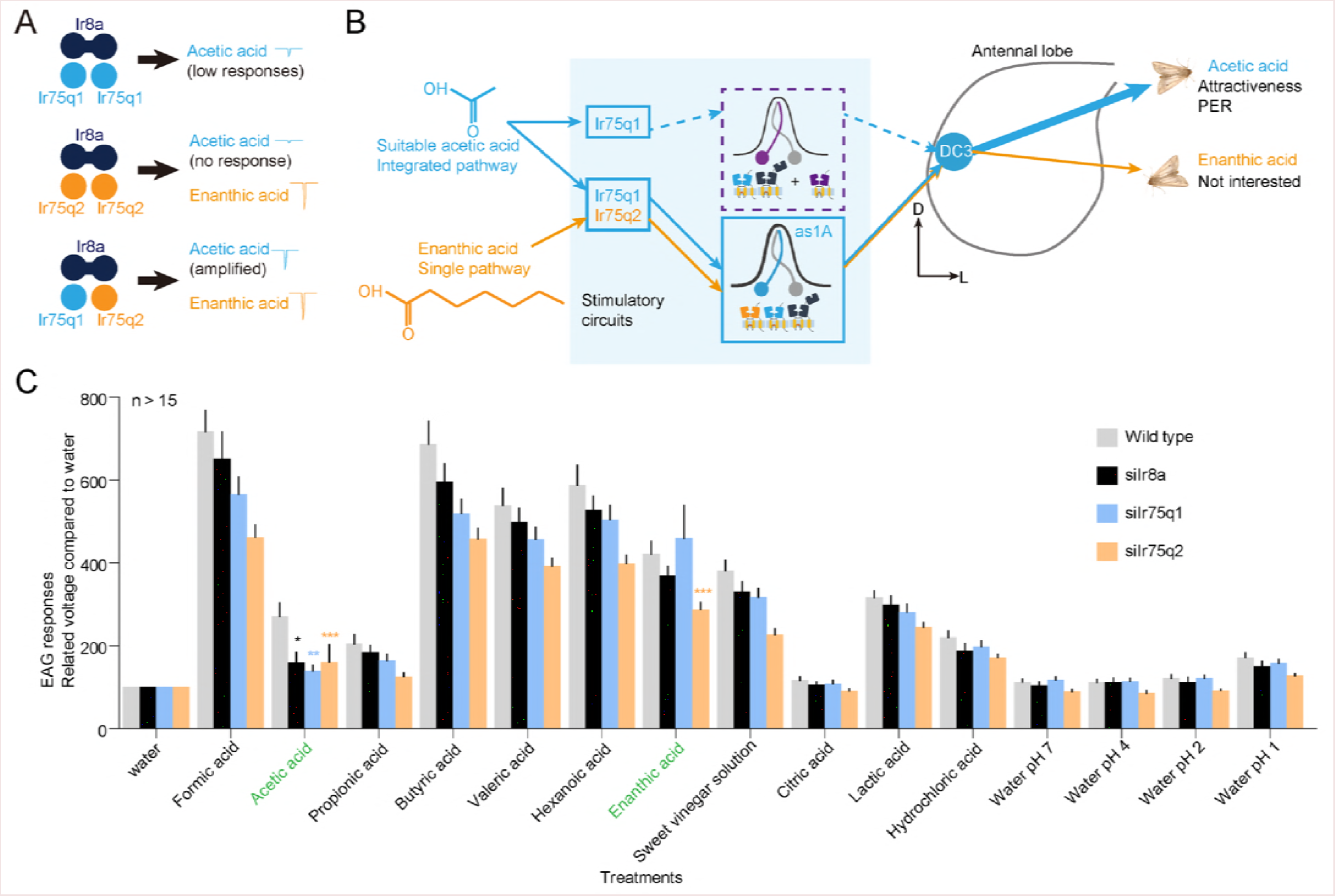
Anticipated molecular basis, coding mechanisms, and behavioral decision of *Ir75q1/q2* mediated olfactory circuits in *M. separata* female moths. **(A)** Schematic diagram showing combinatorial coding patterns of *Ir75q1* and *Ir75q2* according to ionotropic receptor functional structures. Line 1: *Ir75q1* alone delivered acetic acid recognition responses yet at a low reaction rate; line 2: *Ir75q2* alone had an accidental response to enanthic acid yet no preference to acetic acid was shown; line 3: *Ir75q1* + *Ir75q2* had an amplified response to acetic acid yet maintained the accidental response to enanthic acid. **(B)** Sketch map shows olfactory mechanisms of *Ir75q1/q2* pathway at peripheral and antennal lobe level, as well as separation of behavioral consequences between acetic acid and enanthic acid. Enanthic acid activates half of the ORNs (*Ir75q1/q2* expressed) in DC3 pathway, which, does not overcome the threshold to deliver a behavioral output. Suitable amount of acetic acid activates all *Ir75q1* expressed ORNs and they integrate into DC3 to successfully stimulate attractiveness. Dotted lines indicate olfactory pathways for acid avoidance (acids with multi-glomeruli reaction) which were predicted from electrophysiological, behavioral and RNAi test results. **(C)** Electroantennogram assessment of the above model by knocking-down of single gene including *Ir8a* (n = 16), *Ir75q1* (n = 20), and *Ir75q2* (n = 24), respectively. Wild type adults (n = 27) were used as reference. Normalized data were shown with comparing origin voltages of each treatment to water control. Asterisks indicate significantly decrease of electrophysiological responses in sensing acetic acid (Mann Whitney *U* test, siIr8a vs wild type: *P* = 0.013; siIr75q1 vs wild type: *P* = 0.0013; siIr75q2 vs wild type: *P* = 0.0004) or enanthic acid (Mann Whitney *U* test, siIr8a vs wild type: *P* = 0.198; siIr75q1 vs wild type: *P* = 0.296; siIr75q2 vs wild type: *P* = 0.0004) of each treatment comparing to wild type strain. Error bars indicate + s.e.m.

## Discussion

### Combinational coding of *Ir75q1* and *Ir75q2* in *Ir75q1/2*-as-DC3 olfactory circuit

In the current study on *M. separata*, we first proved that acetic acid is the major bio-active compound in the sweet vinegar solution. Then we found 3 types of acid sensing sensilla which had a broad recognition spectrum to multiple acids. Considering so far reported broad spectrum IR-expressed ORNs in *Drosophila* [30], and by comparing amplitudes for all active acids, we suggest that only one ORN was activated for each sensilla type. Although the high responses to enanthic acid of as1A cannot be overlooked, odorant recognizing activities at peripheral level were not always sufficient to deliver final behavioral outputs [49]. Paralleling with this suggestion, we found that acetic acid other than enanthic acid elicited both attractiveness and PER in fasted moths. Similar as it has been reported in *Drosophila* [50,51], it suggests that acetic acid serves as a food related odorant cue and drives initial orientation flights in *M. separata*. Later we located the dedicated glomerulus DC3 in the antennal lobe. The evoking patterns of glomeruli shifted with increasing of acetic acid dosages, reflecting behavioral changes of *M. separata* to acetic acid are determined via these glomeruli [16]. Moreover, the conflict of enanthic acid stimulated reactions between ORN level and antennal lobe level reveal that the behavioral consequences of lower dosage attractiveness has been largely determined via peripheral olfactory coding process. This assumption has been confirmed with later providing evidences on molecular level. According to the published works on IRs, a dimer-dimer complex formed by four monomeric structures (*Ir8a* + *Irx/Irx*) from two genes is needed to perform acid sensing responsiveness [20]. As a result of this, *Ir75q1* + *Ir8a* alone delivered acetic acid recognition responses yet at a low reaction rate, while *Ir75q2* + *Ir8a* alone had an unexpected response to enanthic acid yet no preference to acetic acid was shown. *Ir75q1* + *Ir75q2* + *Ir8a* had an amplified response to acetic acid yet maintained the response to enanthic acid (Figure 8A). The response to enanthic acid could be an evolutionary accident when *M. separata* employ *Ir75q2* to enhance acetic acid sensing in the *Ir75q1* pathway (Figure S8E).

To sum up, the response pattern of oocytes expressing *Ir75q1/q2/8a* correlated with as1A ORNs, and the as1A neuron projected to DC3 together with integration of other sensilla types which can also be reversely proved by RNAi tests. The responses to enanthic acid caused by *Ir75q2* is controlled by lower its expression level, so that more *Ir75q1* can be employed in other sensilla without respond to enanthic acid to provide more acetic acid induced signaling integrating to DC3 glomeruli for delivering attractive behaviors. In all, *Ir75q1* senses acetic acid stimuli and projects to DC3 glomerulus, and half of the signals are amplified by *Ir75q2* (Figure 8B). The similar integration sensilla-to-glomeruli model has also been found in *Drosophila* olfactory receptor based system [52]. With this model, *M. separata* manages to pursue low acetic acid odorant without showing a preference to a non-food related enanthic acid. Taken together, we suggest that the attractive behavioral outputs of the moth stimulated by acetic acid are largely determined via this circuit. It is activated under suitable doses of acetic acid stimuli and mediates behaviors of the insect toward odorant components, which shows a classic chemosensory mechanism, similar to that observed in *Drosophila* in terms of olfaction [16] and gustation [48].

### New features in IR-involved olfactory reception

We revealed a phenomenon that three IRs may form a functional complex. The *Ir75q* clade genes are conserved in many species but not in *Drosophila* [19], so that this phenomenon might also exist in other insect species. In fact, a recently reported amino acid-sensing group of *Ir76b* and *Ir20a* genes also provides evidence for this assumption [22]. However, another work also suggests that *Drosophila Ir76b* might function as a co-receptor gene [53]. To date, there is no solid evidence that more than two IR genes can together form a functional protein complex, but the combinations of IR groups and their functions observed here were somehow much more complex than we expected.

### DC1 and AC1 glomeruli

The *Ir64a*-expressing olfactory pathway is necessary and sufficient for *D. melanogaster* to avoid acid stimuli at 5% [18]. The homologous *Ir64a* gene is also found in *M. separata* (Figure 5A). According to 1 receptor - 1 ORN - 1 glomerulus model [54,55], this receptor may be employed likely by ORNs and glomeruli other than as1A & DC3 related pathway for driving a reversed behavioral consequence to high acidity odorants in this moth. According to RNAi test results, DC1 activity was not contributing to acetic acid attractiveness in *M. separata*. Given the fact that this glomerulus was strongly evoked during application of 5000 μg acetic acid when attractiveness was terminated and antennal lobe reaction pattern was changed dramatically, it is highly likely that DC1 involves in the deterrent olfactory pathway which encodes *Ir64a* (Figure 8B).

It has been reported that *Drosophila Ir75a* responds exclusively to acetic acid and propionic acid [1,30,56], and this reaction pattern coincided with the as2A ORNs which showed high responsiveness to both acetic acid and propionic acid. However, we did not identify a homolog of *Drosophila Ir75a*. The recently reported *Ir75a* pseudogene with a premature termination codon might be the reason for failure to identify *Ir75a* in the *M. separata* transcriptome [30]. To date, no single IR gene has been reported to have sufficient responsiveness to acetic acid alone. However, there might be other IRs which combined with *Ir75q1* to confer stronger acetic acid sensory, as it has been shown in correlations between *Ir75q1/q2* with sensilla types and a slightly decrease of AC1 area during RNAi with siIr75q1. This extra pathway could also be employed by some insect species to override repellent signals from *Ir64a* when exposed to high concentrations of acetic acid solutions, which, ensured a higher acidity tolerance (Figure 8B). Although we could not test this anticipation under the scale of the current study, it is worthwhile to design future investigations on this issue.

### Utilization of *Xenopus laevis* functional demonstration system

The heterologous expression system using *Xenopus* oocytes coupled with double electrode voltage clamp recording is a powerful system for functional investigation of cross membrane ion channels [57]. Moreover, this system has been commonly used in studies involving iGluRs and has the flexibility to build protein complexes using multiple genes [58]. Considering the homology between iGluRs and IRs, we believe that the *Xenopus* system is appropriate for testing functions of *M. separata* IR complexes as has been done for *Drosophila* [29]. However, intact oocytes can be killed when applied with high acidity solutions because the solution pH can be changed by ligand components, showing that there are limits to this system. We managed to minimize this disadvantage by applying acids at low concentrations (10^−6^ mol/L) in order not to change the pH levels. Another thought was to analyze *Ir75q1/q2* with reference to neutralized acid solutions at pH 7.5. In this way, we not only successfully characterized *Ir75q1/q2* in terms of its ligand recognition traits, but also further identified necessary genes for forming the functional ion channel complex.

### Enanthic acid

Enanthic acid stimulates robust responsiveness in as1A, indicating that it is the best ligand for *Ir75q* pathway at peripheral. However, this acid does not evoke higher response of DC3 than acetic acid at antennal lobe level, nor of this acid has attractiveness or PER activity to the moth. As it has been reported in *Drosophila*, non-linearized transmission of olfactory activity exists among peripheral inputs and outputs of other sensation levels due to many architectural variances [52,59–61], the differences of odorant selectivity and preference between ORNs and higher olfactory levels may also exist in *M. separata*. The uneven expression of *Ir75q1* and *Ir75q2* has provided the possible explanation for this phenomenon – although they were co-expressed, more projections of *Ir75q1* were integrated into DC3 glomeruli, resulted in a stronger influence than *Ir75q2*. So that a separation of behavioral outputs of the moth to acetic acid and enanthic acid has been caused. Concisely, behavioral responses of the moth are decided via higher level olfactory system (glomeruli) other than peripheral (ORNs), indicating that acetic acid is a more suitable ligand for *Ir75q* pathway when linking with behavioral outputs in nature. To date, few studies have been conducted on enanthic acid [62,63], and neither of the studies have shown evidence that this component could deliver any behavioral consequences in insects. Thus, we suggest that strong recognitions of this chemical is likely to be an evolutionary side-effect other than a functional feature in foraging for *M. separata*. However, we don’t exclude the possible involvement of enanthic acid in mating or oviposition.

### Acetic acid preference via olfactory circuits

Dose-dependent olfactory preferences towards chemicals are common among animals, including humans, nematodes, and also insects [17]. In insects, IRs are employed by several other sensation pathways which cannot be fulfilled by olfactory or gustatory receptors. On the other, there still be IRs such as *Ir75q1* and *Ir75q2* which are not present in *D. melanogaster* but are conserved in many insect species, including Lepidoptera. Therefore, investigating on these genes and their relevant olfactory pathways in non-model insects is also necessary. As it has been well explained in *Drosophila*, IR based ORNs usually express multiple IRs, and such integrations from same IR expressed ORNs in different sensilla types into the same glomerulus were also found. Revealing the functional architecture of the Ir75q-related polymer complex and its relevant olfactory repertoire has extended our current knowledge of IR family genes. The findings from the present study can also provide insights to further understanding of acid sensing mechanisms across insect species.

## Materials and methods

### Chemical analysis

All chemicals and reagents mentioned in this study can be found in Table S4. Tested sweet vinegar solution samples were made by mixing white wine and vinegar at a 1:3 ratio [33]. Samples were desiccated with calcium chloride drying agent columns and then analyzed with an Agilent Technologies 5973 mass spectrometer coupled with an Agilent Technologies 6890N gas chromatography system (Santa Clara, CA, USA) equipped with a quartz capillary column (HP-5, 30 m x 0.25 mm x 0.25 μm; J&W Scientific, Palo Alto, CA, USA). Volatile compounds were identified by crosschecking with the mass spectrum fragment database (NIST 2.0) with GC/MSD ChemStation (Agilent) and confirmed against standard chemical spectrum patterns. Three replicates were conducted for the sample solution blend.

### Electroantennographic detection

GC-EAD system was used to screen for bioactive chemical compounds following standard protocols as described previously [64]. Antennae were processed by cutting both extremes and immediately mounted with two glass capillary Ag/AgCl electrodes containing Kaissling saline [64] and an identical gas chromatography column was used under the same temperature program as for chemical analysis with the detector at 230 ^o^C. The electrode at the distal end of the antenna was connected via an interface box to a signal acquisition interface board (IDAC; Syntech, Kirchzarten, Germany) connected to a computer. Electroantennogram signals and flame ionization detector responses from the gas chromatography were recorded simultaneously. Bioactive chemicals were identified by crosschecking with GC-MS data. At least 3 replicates were performed for each gender in each species.

Dose response curves were tested for acetic acid using concentration gradient water solutions. Five doses were used as treatments, including 0.1 μg, 1 μg, 10 μg, 100 μg and 1000 μg with water alone used as a blank control. Eleven replicates were tested for females and 18 replicates were tested for males with each treatment.

### Wind tunnel assay

The wind tunnel experiments and data process methods were adopted from a previous study on lepidopterous insects in a dark room [65]. One to 3 day newly emerged female *M. separata* moths were collected and blocked into two groups. One group was fed with enough 10% honey water and another group was fasted for over 12 h before the test. The wind tunnel flying section cubic area was 90 cm × 90 cm × 240 cm. Filtered air was provided through the tunnel inlet via a centrifugal fan at 10 cm/s. The outflowing air was collected by another fan and released through a sealed pipe into the atmosphere. The flight section was lit diffusely from above with red lights at 10 lux. The room temperature was kept at 23 + 2 °C and 40–60% relative humidity. Experiments were conducted at the 4^th^ to 8^th^ hour of the dark phase. Tested chemical solutions were prepared on the day of testing and white filter paper loaded with 10 μl solutions were used as odorant sources. Olfactory stimuli were released from the center of the upwind end of the tunnel and moths were released from a metal mesh cage at the center of the downwind end. Behaviors were recorded for 5 min in terms of taking off, upwind flight, close searching, and landing. Each moth was calculated with an average in each behavior (observed = 1, not observed = 0). Twenty-five (fed) and 35 (fasted) moth adults were tested for each odorant, respectively.

### Proboscis extension response (PER) assay

This assay was adopted from published works in insect [40] (Figure 3E). Tested chemicals or mixtures were loaded with rubber septa with water as control. Naive moths were kept starving in a glass tube (diameter = 2 cm, length = 10 cm) for 12 h and then odorant source tube was connected top-to-top with the insect containing tube. A cotton ball was used in the bottom of the odorant source tube to avoid direct contact of the moths to chemicals. PER reactions were observed within 15 min and 19 individuals were tested for each treatment. Each moth was calculated with an average in PER response (PER observed = 1, not observed = 0). Differences among chemicals were tested with GLM and Tukey HSD multiple comparison tests at *P* = 0.05.

### Single sensillum recording

Two to 3 day old moths were mounted with dental wax inside a 1 ml tip-cut Eppendorf tube and then fixed on a mounting block with the moth’s antennae stretched out sensilla side up. The reference electrode was inserted into a compound eye, and the sharpened recording tungsten electrode was inserted into the base of a single sensillum in the front area of the antenna. Spike numbers in 1 s were calculated by 200 ms spikes timed 5 [66]. Spike sorting and tempo distribution analysis were done referring to previously reported works on moths and vinegar flies [67,68]. Thirty female adults were recorded and among, 3 moths were recorded 5 times in each segment from base to tip of the whole antenna to observe distributions for acid sensilla.

### Antennal lobe calcium imaging

A single 2-3 day old *M. separata* adult was mounted in an artificial mounting block with its antennae being stretched out for chemical stimuli. After dissecting and exposing the brain, the antennal lobe was stained with a calcium-sensitive dye, Calcium Green and Pluronic F-127 mix for one hour at 13 °C, and then thoroughly rinsed with Ringer solution. For imaging, we used a Till Photonics imaging system equipped with a CCD camera connected to an upright microscope. The antennal lobe was illuminated at 475 nm and odorant stimulation started at frame 13 and lasted 500 ms in the recording sequence of 40 frames. The tempo - fluorescence data was transferred and peak 2 to 3 reaction values were selected among frames 13 to 18 [69] for statistics. A total 8 moths were recorded for acid patterns, and 6 moths were recorded for acetic acid dosage responses. Reactions of DC3 glomerulus were cross correlated along 40 frames and the areas during stimulations were analyzed with Pearson’s correlations.

### Antennal lobe atlas

Brains of moths were dissected and processed according to published works [43] and SYNORF1 (Developmental Studies Hybridoma Bank, IA, USA) antibody was used to label glomeruli in antennal lobes. Labelled antennal lobes were visualized with Alexa Fluor 488 goat anti-mouse secondary antibody (Invitrogen) and photo stacks were obtained with a Zeiss LSM710 Meta laser scanning microscope (Zeiss, Oberkochen, Germany). Atlas of *M. separata* brain was done using AMERA 6.0 software (ZIB, Germany). Seven female adults were tested.

### Antennal transcriptome

According to a previous report [45] antennae of male and female *M. separata* moths were collected and stored at −80°C for transcriptome sequencing. Total RNA was extracted using an RNeasy Mini Kit and reverse transcription of cDNA and Illumina library development were performed for Illumina HiSeq2500 sequencing at BGI Co., Beijing, China. High quality clean reads of nuclear sequences were obtained by removing adaptor sequences, empty reads and low-quality sequences (N > 10% sequences) and the reads with more than 50% Q < 20 base using FastQC tool. Clean reads data were combined and *de novo* assembled with Trinity [70]. *Ir* annotations were manually done against reported *Ir* genes with BLAST. Protein structures of genes of interest were predicted with Swiss-model. Translated amino acid sequences were first aligned with ClustalW and phylogenetic tree was developed using the Neighbor-Joining method [71] in MEGA 7.0.14 software [72].

### Quantitative PCR

Reverse transcription PCR was used to analyze tissue expression patterns of related IRs among *M. separata* male antennae (MA), male labial palp (ML), male proboscis (MP), male tarsi (MT), male wing (MW), female antennae (FA), female labial palp (FL), female proboscis (FP), female tarsi (FT), female wing (FW) and ovipositor (O). A total 3 technical replicates were done for RT-PCR tests. Realtime fluorescence quantitative PCR with SYBR Green master mix system (Invitrogen) was used to analyze expression levels of related IRs in RNA interfered *M. separata* strains. Reactions were run with a QuantStudio 3 Real-time PCR system (Applied Biosystems, ThermoFisher Scientific Inc., USA). Gene expression levels were calculated with a formula 1/power (2, Ct *Irx* - Ct *Ir8a*). Primers were listed in Table S2.

### *Xenopus* oocyte expression system and double electrode voltage clamp

Functional analysis was performed as described previously [73]. Full-length cRNA sequences of putative IRs were prepared with mMESSAGE mMACHINE SP6 and injected into *Xenopus laevis* oocytes both individually and in combination. Two-electrode voltage clamp recording was used to detect the whole cell current and oocytes were tested with acids or a series of gradient acetic acid solutions. Intact oocytes were used as control cells for each round of experiment. Data were recorded and processed using pCLAMP Software Suite (Axon Instruments, CA, USA). Treatments included intact cell control, Ir8a, Ir75q1, Ir75q2, Ir8a/Ir75q1, Ir8a/Ir75q2, Ir8a/Ir75q1/Ir75q2, and Ir75q1/Ir75q2, respectively.

### Florescence *in situ* hybridization

Two-color *in situ* hybridization was performed as described previously [45]. Probes of tested IRs were labeled with digoxin or biotin using an RNA labeling Kit version 12 (SP6/T7), with Dig-NTP or Bio-NTP labeling mixture, respectively. Visualization of hybridization signals was performed by successively incubating the sections with HNPP/Fast Red and Biotinyl Tyramide Working Solution with the TSA kit protocol. All sections were analyzed under Zeiss LSM710 microscope. A total 20 pairs of antennae were tested for each treatment.

### RNA interference

siRNAs were prepared with T7 RiboMAX Express RNAi System. Three μg siRNA was injected into a single 3-day pupa of *M. separata*. Moths were then tested after emergence with quantitative PCR (n = 4, with 3 technical replicates each), wind tunnel (n = 18) and electrophysiological tests (n = 9 for SSR and n > 15 for EAG). Non injected strain was used as control, and siGFP strain was used for off-target effect assessment. Treatments included siIr75q1/siIr75q2, siGFP, siIr8a, siIr75q1, and siIr75q2, respectively.

## Data process

All statistics were carried out using IBM SPSS Statistics 22.0.0 (SPSS, Chicago, IL, USA). Bar charts were plotted using Prism 5 for Windows ver. 5.01 (GraphPad software, San Diego, CA, USA). Correlation matrix was developed with Statgraphics Centurion XVII (Statpoint Technologies, Inc., VA, USA). Calcium imaging graphics and *k*-means clustering were processed with MATLAB 7.8.0.347 (The MathWorks, Natick, MA, USA).

## Data viability

Gene deposition information for this study can be found in Table S3. Arithmetic for processing clustering and/or correlation are available upon request.

## Acknowledgments

This work was supported by the Strategic Priority Research Program of the Chinese Academy of Sciences (grant number XDB11010300), the National Natural Science Foundation of China (grant number 31471777 and 31130050), and the National Key Research and Development Program of China (grant number 2016YFC1200600). We thank Prof. Feng Zhang and MARA-CABI Joint Laboratory for Bio-safety, IPP-CAAS for supporting Rui Tang’s research works in IOZ-CAS. We thank Prof. Giovanni Galizia for helping with optical imaging data analysis. We thank Dr. Stefan Toepfer and Prof. Fang-Hao Wan for the valuable comments on the manuscript. We thank Rui Wang, Xiao-Wei Qin and Ling Yang from the State Key Laboratory for their assistance with chemical analysis and *in situ* hybridization; Dr. Meng Xu, Dr. Wu Han, Yan Chen and Ke Yang for technical assistance in the SSR, calcium imaging, and molecular biology experiments, respectively. We also thank Dr. Jin-Ping Zhang, Dr. Ya-Lan Sun and Dr. Xiao-Jing Jiang for providing insects for GC-EAD tests, B. F. A. Xiao-Qian Bao for technical support with insect graph development, Dr. Yi-Nan Fang for the technical support with the Matlab clustering analysis.

## Author contributions

CZW conceived the project. RT and CZW designed the study. RT conducted electrophysiological and molecular tests. RT, LQH, and NJJ conducted bioassays. NJJ, CN, and RT conducted antennae transcriptome sequencing and data analysis. RT and CZW analyzed data and wrote the manuscript.

## Experimental procedures

All the live animal procedures used in this study were strictly conducted under guidelines developed by the ethics committee of the State Key Laboratory of Integrated Management of Pest Insects and Rodents (ethic no. IOZ17090-A), Institute of Zoology, Chinese Academy of Sciences. Animal procedures were performed under anesthesia, and wounds were carefully treated to avoid infection. Any suffering was minimized.

## Supporting Information Legends

**Table S1. Results of gas chromatography coupled mass spectrum analysis for chemical fingerprints information in sweet vinegar solution volatiles.** Asterisks indicate bioactive compounds which stimulated electric peaks of female *M. separata* antennae in physiological test.

**Table S2. List of primers used in the study.** F: forward strand; R: reverse strand. Underlined letters indicate restriction recognition sites. Bold letters indicate *Kozak* sequence.

**Table S3. Gene deposition information in this study.**

**Table S4. Chemicals and reagents used in the study.**

**Figure S1.** Electroantennogram responses of nine insect species. **(A)** Gas chromatography - coupled electroantennographic detection (GC-EAD) tests of the sweet vinegar solution with five representative insect species. Obvious amplitudes in the GC-EAD output indicate responses of insect antennae to isolated acetic acid volatile component. **(B)** Comparison of multiple species in terms of GC-EAD responsiveness to acetic acid. Different lower case letters indicate significant differences of electrical activity in the antennae (GLM and Tukey HSD, F_8, 67_ = 7.6, *P* < 0.0001). Error bars indicate + s.e.m.

**Figure S2.** Distribution of acid sensing sensilla on antennae of *M. seprarata* females. **(A)** Distribution of sensilla which responded to acetic acid on female *M. separata* antenna. **(B)** Distributional map of spike counts for three types of sensilla to enanthic acid.

**Figure S3.** Overall assessment of female *M. separata* adults in wind tunnel tests. For each data point n ≥ 10. Benzaldehyde was used as a positive control. ***: significant differences between feeding status within each behavior (GLM, Take off: F_317_ = 23.4, *P* < 0.0001; Upwind flight: F_317_ = 30.4, *P* < 0.0001; Close search: F_317_ = 16.9, *P* < 0.0001). #: significant differences of treatments compared with water control (GLM and Dunnett test referring to water control, acetic acid 10 μg: *P* = 0.02; acetic acid 100 μg: *P* = 0.021; acetic acid 1000 μg: *P* < 0.0001).

**Figure S4.** Antennal lobe optical imaging tests for acids toward female *M. separata* adults. **(A)** Representative reaction patterns of female *M. separata* antennal lobes toward acids. ROI: regions of interest. X-axis: frames. Y-axis: ΔF/F0. Water was used as control. **(B)** Reaction patterns of female *M. separata* antennal lobes toward gradient dosages of acetic acid.

**Figure S5.** Representative electrophoresis result of tissue expression of 4 *Irs* by reverse transcription PCR (RT-PCR) using *M. separata* cDNAs from different body parts. *β*-actin was used as reference. Sampled organs were sketched from left to right as: antennae, labial palps, proboscis, forelegs, wings of males and females, and female ovipositors.

**Figure S6.** Sequence alignment of IRs. **(A)** Sequence alignment of *Ir8a* from *Drosophila melanogaster* and *Mythimna separata*. **(B)** Sequence alignment of *Ir75x* from *Drosophila melanogaster* and *Mythimna separata*. Gene deposition information and ID number were listed in Table S3. ✶ indicate predicted conserved amino acid residues which are essential for ligand recognitions within binding domain of each IR.

**Figure S7.** Structural prediction of 4 IRs from *M. separata*. The structures of IRs were developed via SWISS-MODEL website programme (www.swissmodel.expasy.org). *D. melanogaster* IRs were used as references.

**Figure S8.** Functional analysis of IRs using *Xenopus* oocyte expression system. **(A-E)** All control groups of cells tested with multiple acid solutions. 1C: formic acid, 2C: acetic acid, 3C: propionic acid, 4C: butyric acid, 5C: valeric acid, 6C: hexanoic acid, 7C: enanthic acid. Bars with different lower case letters indicate significant differences of responses caused among tested acids (GLM and Tukey HSD, *Ir75q1/Ir75q2:* F_9, 106_ = 10.8, *P* < 0.0001. Error bars indicate + s.e.m.). **(F)** Comparison of all tested combinations to acetic acid solutions. * and **: significant higher currents elicited by acetic acid. (GLM and Dunnett test against intact oocyte control: F_6, 56_ = 6.6, *P* < 0.0001). NS: no different was observed for *Ir75q1/q2/8a* between 1 mmol/L and 0.001 mmol/L acetic acid stimuli (Student’s *t* test, *t* = 0.637, *P* = 0.534). **(G)** Representative responses of intact control oocytes.

**Figure S9.** Representative responses **(A)** and statistics **(B)** of acidity responses of *Ir8a/Ir75q1/Ir75q2* to normal and neutralized acetic acid solutions with the same concentration of 10^−3^ mol/L. Different letters indicate significant differences of currents among acids (GLM and Tukey HSD, Origin solution: F_5, 45_ = 16.2, *P* < 0.0001; Neutralized solution: F_5, 31_ = 5.4, *P* = 0.001. Error bars indicate + s.e.m.).

**Movie S1.** Close feeding attempt of fasted *M. separata* female adults driven by acetic acid odorant stimuli. Direct contact with acetic acid source was avoided by the cotton ball in the tube to remove possible gustatory reception in the test.

**Movie S2.** Antennal lobe evoking pattern to 5000 μg acetic acid in female *M. separata* adults. Three distinct areas of interests (ROIs) were activated during stimuli at frame 13-18. The three ROIs were named according to the positions as DC1 (up, dorsal central), DC3 (mid), and AC1 (down, arterial central).

**Movie S3.** Antennal lobe evoking pattern to 5000 μg enanthic acid in female *M. separata* adults. Only DC3 glomerulus was activated during stimuli at frame 13-18.

**Movie S4.** Antennal lobe evoking pattern to 5000 μg acetic acid in female *M. separata* adults from siIr75q1/q2 strain. DC1 and AC1 glomeruli were activated during stimuli at frame 13-18, while the area of DC3 was not evoked with an observable reaction.

**Data S1.** Compiled results from SSR experiments of wild type strain. Spikes recorded to water control, acetic acid, and enanthic acid were shown.

**Data S2.** Compiled results from SSR experiments of RNAi strain. Spikes recorded to water control, acetic acid, and enanthic acid were shown.

## References

1. Joseph RM & Carlson JR (2015) Drosophila chemoreceptors: A molecular interface between the chemical world and the brain. Trend Genet : TIG 31(12):683–695.

2. Allmann S, et al. (2013) Feeding-induced rearrangement of green leaf volatiles reduces moth oviposition. Elife 2:e00421.

3. Dweck HK, et al. (2015) Pheromones mediating copulation and attraction in Drosophila. Proc Natl Acad Sci U S A 112(21):E2829–2835.

4. Ebrahim SA, et al. (2015) Drosophila avoids parasitoids by sensing their semiochemicals via a dedicated olfactory circuit. PLoS Biol 13(12):e1002318.

5. Penniston KL, Nakada SY, Holmes RP, & Assimos DG (2008) Quantitative assessment of citric acid in lemon juice, lime juice, and commercially-available fruit juice products. J Endourol 22(3):567–570.

6. Zhu J, Park K-C, & Baker TC (2003) Identification of odors from overripe mango that attract vinegar flies, Drosophila melanogaster. J Chem Ecol 29(4):899–909.

7. Gorter JA, et al. (2016) The nutritional and hedonic value of food modulate sexual receptivity in Drosophila melanogaster females. Sci Rep 6:19441.

8. Gou B, Liu Y, Guntur AR, Stern U, & Yang CH (2014) Mechanosensitive neurons on the internal reproductive tract contribute to egg-laying-induced acetic acid attraction in Drosophila. Cell Rep 9(2):522–530.

9. Joseph RM, Devineni AV, King IF, & Heberlein U, (2009) Oviposition preference for and positional avoidance of acetic acid provide a model for competing behavioral drives in Drosophila. Proc Natl Acad Sci U S A 106(27):11352–11357.

10. Chen Y & Amrein H (2017) Ionotropic receptors mediate drosophila oviposition preference through sour gustatory receptor neurons. Curr Biol27:1–10.

11. Cha DH, Adams T, Rogg H, & Landolt PJ (2012) Identification and field evaluation of fermentation volatiles from wine and vinegar that mediate attraction of spotted wing Drosophila, Drosophila suzukii. J Chem Ecol 38(11):1419–1431.

12. Landolt P & Zhang Q-H (2016) Discovery and development of chemical attractants used to trap pestiferous social wasps (Hymenoptera: Vespidae). J Chem Ecol 42(7):655–665.

13. Landolt PJ & Higbee BS (2002) Both sexes of the true armyworm (Lepidoptera: Noctuidae) trapped with the feeding attractant composed of acetic acid and 3-methyl-1-butanol. Fla Entomol 85(1):182–185.

14. Meagher RL & Mislevy P (2005) Trapping Mocis spp. (Lepidoptera : Noctuidae) adults with different attractants. Fla Entomol 88(4):424–430.

15. Toth M, et al. (2010) Male and female noctuid moths attracted to synthetic lures in Europe. J Chem Ecol 36(6):592–598.

16. Semmelhack JL & Wang JW (2009) Select Drosophila glomeruli mediate innate olfactory attraction and aversion. Nature 459(7244):218–223.

17. Yoshida K, et al. (2012) Odour concentration-dependent olfactory preference change in C. elegans. Nat Comm 3:739.

18. Ai M, et al. (2010) Acid sensing by the Drosophila olfactory system. Nature 468(7324):691–695.

19. Benton R, Vannice KS, Gomez-Diaz C, & Vosshall LB (2009) Variant ionotropic glutamate receptors as chemosensory receptors in Drosophila. Cell 136(1):149–162.

20. Ai M, et al. (2013) Ionotropic glutamate receptors IR64a and IR8a form a functional odorant receptor complex in vivo in Drosophila. JNeuroci 33(26):10741–10749.

21. Prieto-Godino LL, et al. (2017) Evolution of acid-sensing olfactory circuits in Drosophilids. Neuron 93(3):661–676.e666.

22. Ganguly A, et al. (2017) A molecular and cellular context-dependent role for Ir76b in detection of amino acid taste. Cell Rep 18(3):737–750.

23. Hussain A, et al. (2016) Ionotropic chemosensory receptors mediate the taste and smell of polyamines. PLoS Biol 14(5):e1002454.

24. Koh TW, et al. (2014) The Drosophila IR20a clade of ionotropic receptors are candidate taste and pheromone receptors. Neuron 83(4):850–865.

25. Lee Y, Poudel S, Kim Y, Thakur D, & Montell C (2018) Calcium taste avoidance in Drosophila. Neuron 97(1):67–74 e64.

26. Ni L, et al. (2016) The ionotropic receptors IR21a and IR25a mediate cool sensing in Drosophila. Elife e13254.

27. Enjin A, et al. (2016) Humidity sensing in Drosophila. Curr Biol 26(10):1352–1358.

28. Knecht ZA, et al. (2016) Distinct combinations of variant ionotropic glutamate receptors mediate thermosensation and hygrosensation in Drosophila. Elifee17879.

29. Abuin L, et al. (2011) Functional architecture of olfactory ionotropic glutamate receptors. Neuron 69(1):44–60.

30. Prieto-Godino LL, et al. (2016) Olfactory receptor pseudo-pseudogenes. Nature 539(7627): 93–97.

31. Bisch-Knaden S, Dahake A, Sachse S, Knaden M, & Hansson BS (2018) Spatial representation of feeding and oviposition odors in the brain of a hawkmoth. Cell Rep 22(9):2482–2492.

32. Faucher CP, Hilker M, & de Bruyne M (2013) Interactions of carbon dioxide and food odours in Drosophila, olfactory hedonics and sensory neuron properties. PloS One 8(2):e56361.

33. Jiang XF, Zhang L, Cheng YX, & Luo LZ (2014) Current status and trends in research on the oriental armyworm, Mythimna separata (Walker) in China. Chin J Appl Entomol 51(4):881–889.

34. Baini F, Del Vecchio M, Vizzari L, & Zapparoli M (2016) Can the efficiency of pitfall traps in collecting arthropods vary according to the used mixtures as bait? Rendiconti Lincei 27(3):495–499.

35. Landolt PJ, Smithhisler CS, Reed HC, & McDonough LM (2000) Trapping social wasps (Hymenoptera: Vespidae) with acetic acid and saturated short chain alcohols. J Econom Entomol 93(6):1613–1618.

36. Olivier V, Monsempes C, Francois MC, Poivet E, & Jacquin-Joly E (2011) Candidate chemosensory ionotropic receptors in a Lepidoptera. insect Mol Biol 20(2):189–199.

37. van Schooten B, Jiggins CD, Briscoe AD, & Papa R (2016) Genome-wide analysis of ionotropic receptors provides insight into their evolution in Heliconius butterflies. BMC Genom 17(1):254.

38. Macqueen J (1967) Some methods for classification and analysis of multivariate observations. Proc. of Berkeley Symposium on Mathematical Statistics and Probability, pp 281–297.

39. Rousseeuw PJ (1987) Silhouettes: A graphical aid to the interpretation and validation of cluster analysis. J Comp Appl Math 20(20):53–65.

40. Smith BH & Burden CM (2014) A proboscis extension response protocol for investigating behavioral plasticity in insects: Application to basic, biomedical, and agricultural research. J Vis Exp (91).

41. Berg BG, Galizia CG, Brandt R, & Mustaparta H (2002) Digital atlases of the antennal lobe in two species of tobacco budworm moths, the Oriental Helicoverpa assulta (male) and the American Heliothis virescens (male and female). J Comp Neurol 446(2):123–134.

42. Lofaldli BB, Kvello P, & Mustaparta H (2010) Integration of the antennal lobe glomeruli and three projection neurons in the standard brain atlas of the moth Heliothis virescens. virescens. Front Sys Neurosci 4:5.

43. Zhao XC, et al. (2016) Glomerular identification in the antennal lobe of the male moth Helicoverpa armigera. J Comp Neurol 524(15):2993–3013.

44. Yang K, Huang LQ, Ning C, & Wang CZ (2017) Two single-point mutations shift the ligand selectivity of a pheromone receptor between two closely related moth species. Elife 6.

45. Ning C, Yang K, Xu M, Huang LQ, & Wang CZ (2016) Functional validation of the carbon dioxide receptor in labial palps of Helicoverpa armigera moths. insect Biochem Mol Biol 73:12–19.

46. Clark JT & Ray A (2016) Olfactory mechanisms for discovery of odorants to reduce insect-host contact. J Chem Ecol 42(9):919–930.

47. Strutz A, et al. (2014) Decoding odor quality and intensity in the Drosophila brain. Elife 3:e04147.

48. Zhang YV, Ni J, & Montell C (2013) The molecular basis for attractive salt-taste coding in Drosophila. Science 340(6138):1334–1338.

49. Choo YM, et al. (2018) Reverse chemical ecology approach for the identification of an oviposition attractant for Culex quinquefasciatus. Proc Natl Acad Sci U S A 115(4):714–719.

50. Ko K, et al. (2015) Starvation promotes concerted modulation of appetitive olfactory behavior via parallel neuromodulatory circuits. Elife 4:e08298.

51. Lebreton S, Becher PG, Hansson BS, & Witzgall P (2012) Attraction of Drosophila melanogaster males to food-related and fly odours. J Insect Physiol 58(1):125–129.

52. Grabe V, et al. (2016) Elucidating the neuronal architecture of olfactory glomeruli in the Drosophila antennal lobe. Cell Rep 16(12):3401–3413.

53. Croset V, Schleyer M, Arguello JR, Gerber B, & Benton R (2016) A molecular and neuronal basis for amino acid sensing in the Drosophila larva. Sci Rep 6:34871.

54. Galizia CG, Munch D, Strauch M, Nissler A, & Ma S (2010) Integrating heterogeneous odor response data into a common response model: A DoOR to the complete olfactome. Chem Sens 35(7):551–563.

55. Vosshall LB & Stocker RF (2007) Molecular architecture of smell and taste in Drosophila. Annu Rev Neurosci 30:505–533.

56. Rytz R, Croset V, & Benton R (2013) Ionotropic receptors (IRs): chemosensory ionotropic glutamate receptors in Drosophila and beyond. Insect Biochem Mol Biol 43(9):888–897.

57. Dascal N (2008) The use of Xenopus oocytes for the study of ion channel. Crit Rev Biochem 22(4):317–387.

58. Walker CS, et al. (2006) Reconstitution of invertebrate glutamate receptor function depends on stargazin-like proteins. Proc Natl Acad Sci U S A 103(28):10781–10786.

59. Bhandawat V, Olsen SR, Gouwens NW, Schlief ML, & Wilson RI (2007) Sensory processing in the Drosophila antennal lobe increases reliability and separability of ensemble odor representations. Nat Neurosci 10(11):1474–1482.

60. Caron SJ, Ruta V, Abbott LF, & Axel R (2013) Random convergence of olfactory inputs in the Drosophila mushroom body. Nature 497(7447):113–117.

61. Chou YH, et al. (2010) Diversity and wiring variability of olfactory local interneurons in the Drosophila antennal lobe. Nat Neurosci 13(4):439–449.

62. Urbanek A, et al. (2012) Composition and antimicrobial activity of fatty acids detected in the hygroscopic secretion collected from the secretory setae of larvae of the biting midge Forcipomyia nigra (Diptera: Ceratopogonidae). J Insect Physiol 58(9):1265–1276.

63. Watanabe H, Ai H, & Yokohari F (2012) Spatio-temporal activity patterns of odor-induced synchronized potentials revealed by voltage-sensitive dye imaging and intracellular recording in the antennal lobe of the cockroach. Front Sys Neurosci 6(55):1–18.

64. Tang R, et al. (2016) Identification and testing of oviposition attractant chemical compounds for Musca domestica. Sci Rep 6,33017.

65. Tang R, Zhang JP, & Zhang ZN (2012) Electrophysiological and behavioral responses of male fall webworm moths (Hyphantria cunea) to herbivory-induced mulberry (Morus alba) leaf volatiles. PloS One 7(11):e49256.

66. Xu M, et al. (2016) Olfactory perception and behavioral effects of sex pheromone gland components in Helicoverpa armigera and Helicoverpa assulta. Sci Rep 6:22998.

67. de Bruyne M, Foster K, & Carlson JR (2001) Odor coding in the Drosophila antenna. Neuron 30(2):537–552.

68. Ghaninia M, Olsson SB, & Hansson BS (2014) Physiological organization and topographic mapping of the antennal olfactory sensory neurons in female hawkmoths, Manduca sexta. Chem Sens 39:655–671.

69. Wu H, et al. (2015) Specific olfactory neurons and glomeruli are associated to differences in behavioral responses to pheromone components between two Helicoverpa species. Front Behav Neurosci 9:206.

70. Haas BJ, et al. (2013) De novo transcript sequence reconstruction from RNA-seq using the Trinity platform for reference generation and analysis. Nat Prot 8(8):1494–1512.

71. Saitou N & Nei M (1987) The Neighbor-joining method: a new method for reconstructing phylogenetic trees. Mol Biol Evol 4(4):406–425.

72. Kumar S, Stecher G, & Tamura K (2016) MEGA7: Molecular Evolutionary Genetics Analysis version 7.0 for bigger datasets. Mol Biol Evol 33(7):1870–1874.

73. Di C, Ning C, Huang LQ, & Wang CZ (2017) Design of larval chemical attractants based on odorant response spectra of odorant receptors in the cotton bollworm. Insect Biochem Mol Biol 84:48–62.

